# Visual adaptation of opsin gene expression to the aquatic environment in sea turtles

**DOI:** 10.1101/2023.08.24.554587

**Authors:** Yohey Terai, Misa Osada, Satomi Kondo, Masayoshi Tokita

## Abstract

Several vertebrate taxa, cetaceans, sirenia, pinnipeds, and sea snakes have adapted to the marine aquatic environment. In the species of these taxa, marine adaptation has resulted in shifts in the absorption spectra of opsin pigments and/or the degeneration or duplication of opsin genes. Thus, marine adaptation has strongly affected the evolution of opsins. In sea turtles, however, the effect of adaptation from freshwater to marine environments on opsin evolution has not been studied. In this study, we determined the high-throughput RNA sequences extracted from eyes of two sea turtles (green turtle: *Chelonia mydas*, loggerhead: *Caretta caretta*) and two freshwater turtle species (three-keeled pond turtle: *Mauremys reevesii*, softshell turtle: *Pelodiscus sinensis*) and investigated the amino acid evolution and expression of the opsin gene. We found that most of the sea turtle lineage-specific amino acid substitutions did not alter amino acid properties and did not include previously known substitutions for turning absorption spectra of opsin pigments, suggesting no adaptive amino acid substitutions in the opsins during marine adaptation in sea turtles. Instead, the blue-sensitive opsin (*SWS2*) gene expression was higher in sea turtles than in freshwater turtles. These results suggest that sea turtles may have adapted their vision to the blue light-rich marine environment by increasing *SWS2* expression.

## Introduction

Several taxa of terrestrial vertebrates have returned to the sea and adapted to the aquatic environment. The cetaceans, sirenia, and the hydrophiin sea snakes are fully adapted to aquatic environments and no longer depend on terrestrial habitats (Heatwole 1999; Sanders and Lee 2010). These three groups of species are entirely aquatic, giving birth to their young in the water and spending their entire lives in the water. Some vertebrate taxa spend part of their life above water: pinnipeds and amphibious laticaudins sea snakes come ashore to spawn, breed, and rest but forages in the water. Sea turtles spend most of their lives underwater but lay their eggs on land.

Sea turtles (Chelonioidea superfamily) are an ancient diverged species (Bowen and Karl 2007) that originated in the Late Jurassic (Naro-Maciel, et al. 2008; Joyce, et al. 2013). Seven sea turtle species are currently recognized, with loggerhead, green, and leatherback turtles found in tropical and subtropical waters worldwide. Molecular phylogenetic analysis indicates that sea turtles form a monophyletic group, which is sister to the freshwater turtles (Thomson, et al. 2021). In other words, sea turtles likely adapted from freshwater to a marine environment.

Visual adaptation is a prime example of molecular evolution by natural selection (Davies, et al. 2012). Vertebrate visual pigments comprise a light-absorbing component (chromophore) and a protein component called opsin (Shichida 1999). The spectral sensitivity of a photoreceptor is determined by the chromophore type and the chromophore’s interaction with the amino acid residues in the retinal binding pocket where the chromophore lies (Yokoyama 2000). Several spectral tuning positions in the amino acid sequence of opsins have been reported, and these positions have been used to predict the absorption spectra of visual pigments from opsin gene sequences (Yokoyama and Radlwimmer 1998, 1999; Yokoyama 2000; Hunt, et al. 2001; Yokoyama 2008; Yokoyama, et al. 2008). Vertebrates have two types of photoreceptor cells that differ in structure and function: cone cells, which are active in bright light and contain cone visual pigments (SWS1, SWS2, RH2, LWS) for color vision, and rod cells, which contain RH1 pigments for scotopic vision (Yokoyama 2000; Bowmaker 2008; Davies, et al. 2012).

The transition from terrestrial to marine environments has affected the evolution of opsins in marine mammals and sea snakes. A correlation between foraging depth and spectral sensitivity of visual pigments has been reported in cetaceans (McFarland 1971; Fasick, et al. 1998; Fasick and Robinson 1998; Fasick and Robinson 2000; Bischoff, et al. 2012). Shallow-diving species have a peak absorbance (λmax) of RH1 comparable to that of land mammals (500 nm), while deep-diving species have a blue shift in λmax around 479 nm (Fasick and Robinson 2000; Bischoff, et al. 2012). A correlation between foraging depth and the spectral sensitivity of RH1 pigments has also been reported in pinnipeds. Deep-diving elephant seals have a λmax of 483 nm, while other shallow-diving pinnipeds have a λmax similar to land mammals (Southall, et al. 2002; Levenson, et al. 2006). A correlation between foraging depth and spectral sensitivity of visual pigments and the amino acid substitutions responsible for this has also been reported in sea snakes (Seiko, et al. 2020). The blueshifts of LWS pigments have occurred stepwise and may have adapted to the deep-sea or open-ocean environment where blue light is dominant (Seiko, et al. 2020). In Hydrophiini, a substitution causes a long wavelength shift in the SWS1 pigment, and the polymorphism at this position has persists among Hydrophiini species (Simões, et al. 2020).

In addition to spectral shifts, opsin genes have been lost or duplicated in marine mammals and sea snakes during adaptation to the marine environment. All cetaceans have lost the SWS1 gene (Fasick, et al. 1998; Peichl, et al. 2001; Griebel and Peichl 2003; Levenson and Dizon 2003) at an early stage of evolutionary transition (Meredith, et al. 2013). All pinnipeds also lost the SWS1 gene (Peichl and Moutairou 1998; Peichl, et al. 2001; Newman and Robinson 2005; Levenson, et al. 2006), indicating that cetaceans and pinnipeds convergently degenerated *SWS1* gene during the evolutionary transition. Furthermore, cetaceans have independently lost LWS multiple times in both toothed and baleen whale lineages (Meredith, et al. 2013). No gene loss has occurred in sea snakes, while a possible duplication of the *SWS1* gene has been suggested (Rossetto, et al. 2023).

Marine adaptation has strongly affected the evolution of opsin in cetaceans, pinnipeds, and sea snakes. In sea turtles, however, the effect of adaptation from freshwater to marine environments on opsin evolution has not been studied. In this study, we examined the amino acid evolution and expression of the opsin gene and showed the effects of marine adaptation on the opsin gene in sea turtles.

## Results

### Phylogenetic relationships of opsin gene sequences

We used these sequences of loggerhead turtle *SWS1, SWS2, RH2, LWS*, and *RH*1 sequences in the database (Table S1). Using these opsin sequences as queries, we searched the opsin sequences of Testudines species from the database by blastn (Table S1). We also used the most similar outgroup sequences as outgroup sequences for phylogenetic analysis (Table S1). Some opsin gene sequences were not in the database for green and softshell turtles. Therefore, we assembled eye-derived RNA-seq reads and isolated opsin sequences from each assembled sequence by blastn using the loggerhead turtle opsin sequence as queries (Table S1). We then constructed phylogenetic trees from the sequences of *SWS1, SWS2, RH2, LWS*, and *RH1*, respectively, using the ML method (Fig. 1). As for the phylogenetic relationships among turtle species, the topologies were the same as those of previous studies (Thomson, et al. 2021), except for SWS2.

**Figure 1.**
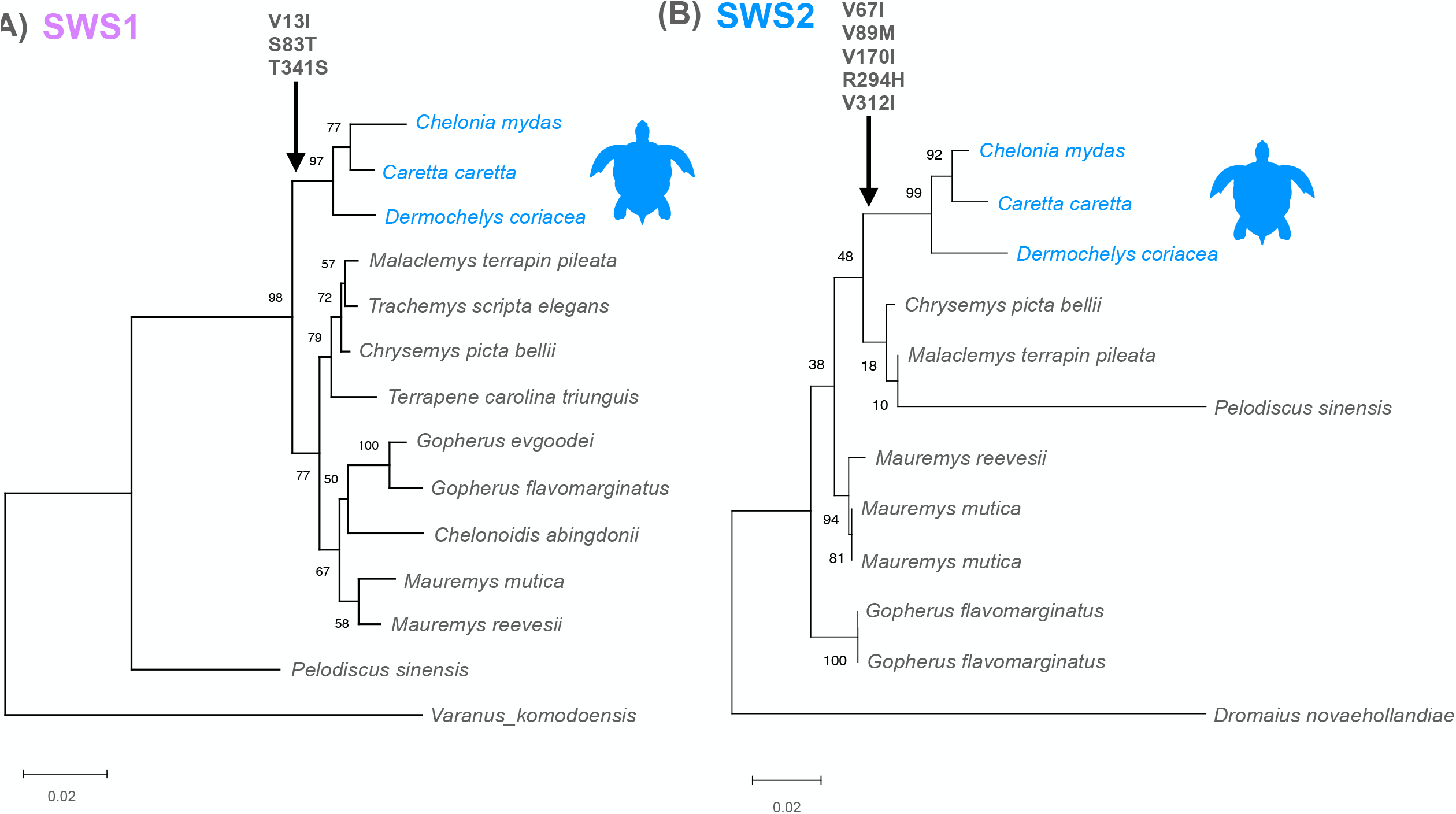
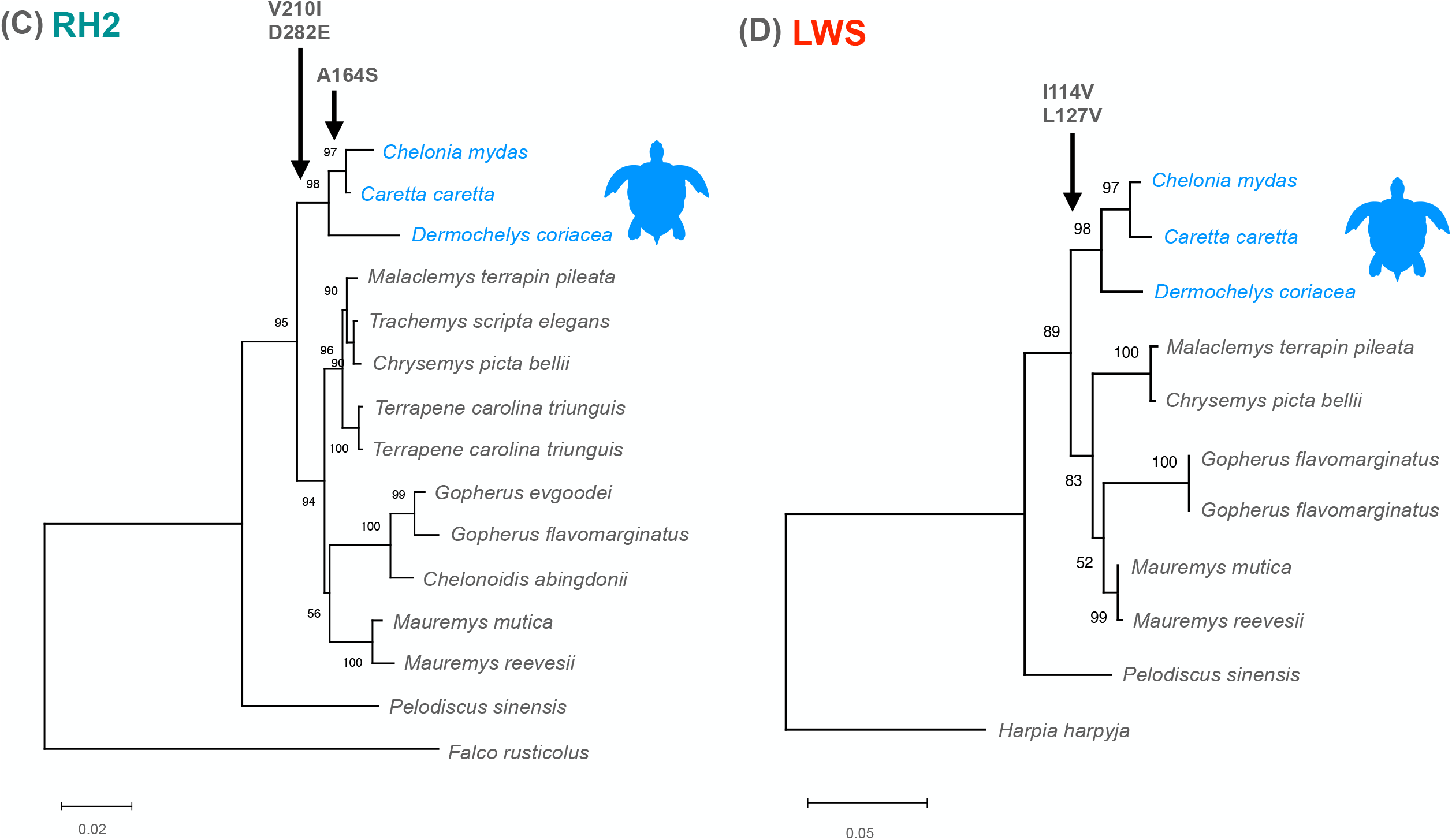
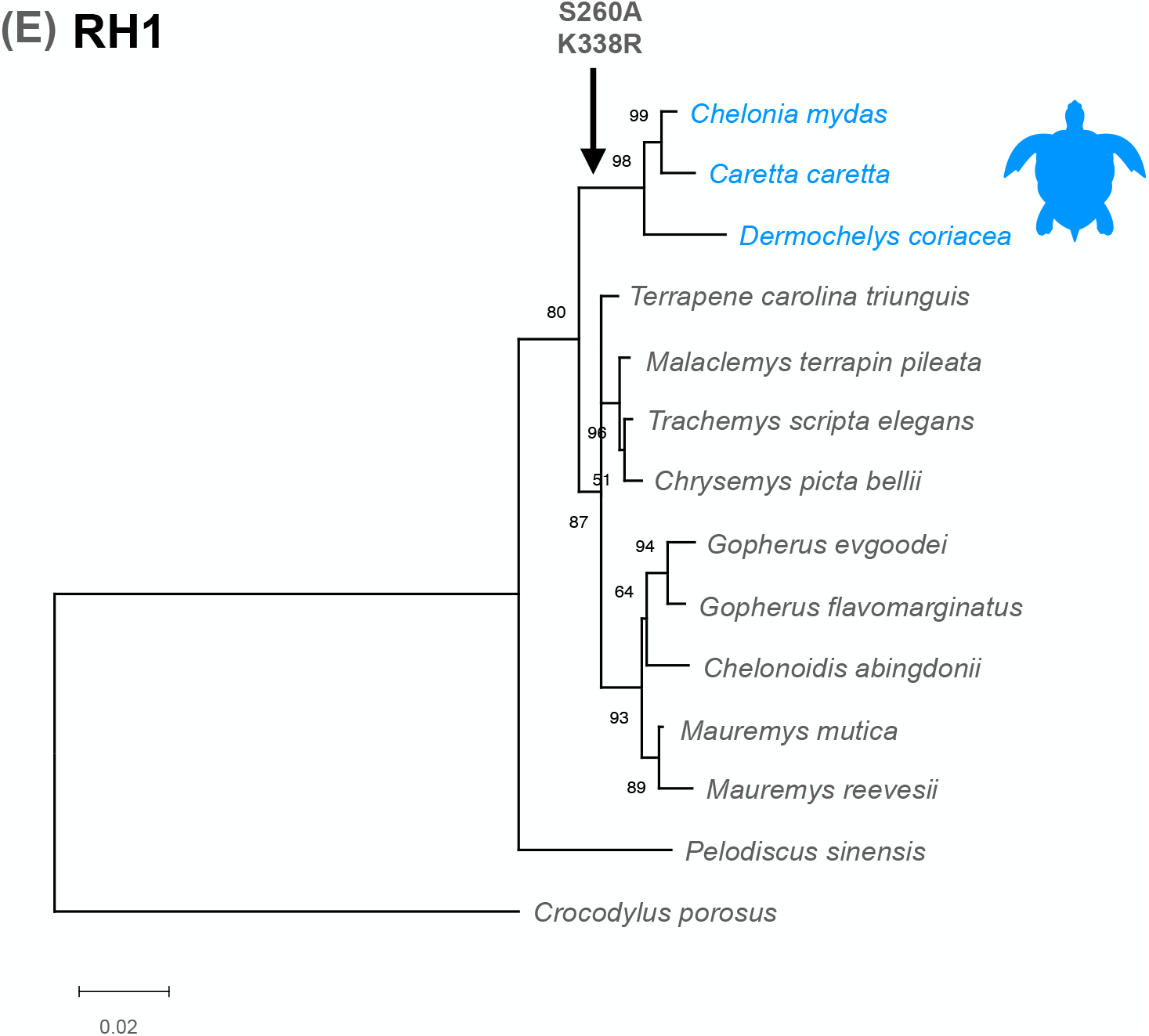
Muximum likelihood trees of (A) SWS1, (B) SWS2, (C) RH2, (D) LWS, and (E) RH1. The arrow indicates the position of the common ancestor of the sea turtles with the amino acid substitution specific to sea turtle lineage.

### Sea turtle lineage-specific amino acid substitutions

To search amino acid substitutions that played an adaptive role in the adaptation of sea turtles to the marine environment, we estimated and placed sea turtle lineage-specific amino acid substitutions on the phylogenetic trees of the respective opsin genes (Fig. 1). Most of these sea turtle lineage-specific amino acid substitutions did not alter amino acid properties. In addition, the sea turtle lineage-specific substitutions did not include previously known substitutions for turning absorption spectra of opsin pigments. These results suggest that no adaptive amino acid substitutions may have occurred in the opsins during marine adaptation in sea turtles. The only amino acid substitution in the RH2, position 164, substituted from alanine to serine in green and loggerhead turtles, was estimated to be a long-wavelength shift of the absorption about 7 nm (Yokoyama and Radlwimmer 1998, 1999; Yokoyama 2000; Hunt, et al. 2001; Yokoyama 2008; Yokoyama, et al. 2008). This substitution may be related to the ecology of green and loggerhead turtles but not to marine adaptation since it is alanine in leatherback turtle.

### Expression patterns of opsin genes

To examine opsin gene expression patterns in green, loggerhead, three-keeled pond, and softshell turtles, RNA-seq reads from each species’ eyes were mapped to each species’ opsin gene. We calculated the TPM (Transcript Per Million) from the mapping results. Then we calculated the relative expression levels of the opsin genes (Fig. 2). The expression patterns of two sea turtle species (Figs. 2A and 2B) and two freshwater turtle species (Figs. 2C and 2D) were similar, except for *SWS2. SWS2* expression tended to be higher in the two sea turtle species. Indeed, we compared the expression levels of each opsin gene between sea turtles and freshwater turtles (Fig. 3) and found that only *SWS2* was significantly higher in sea turtles than in freshwater turtles (Fig. 3B). These results suggest that sensitivity to blue light may be higher in sea turtles than in freshwater turtles.

**Figure 2.**
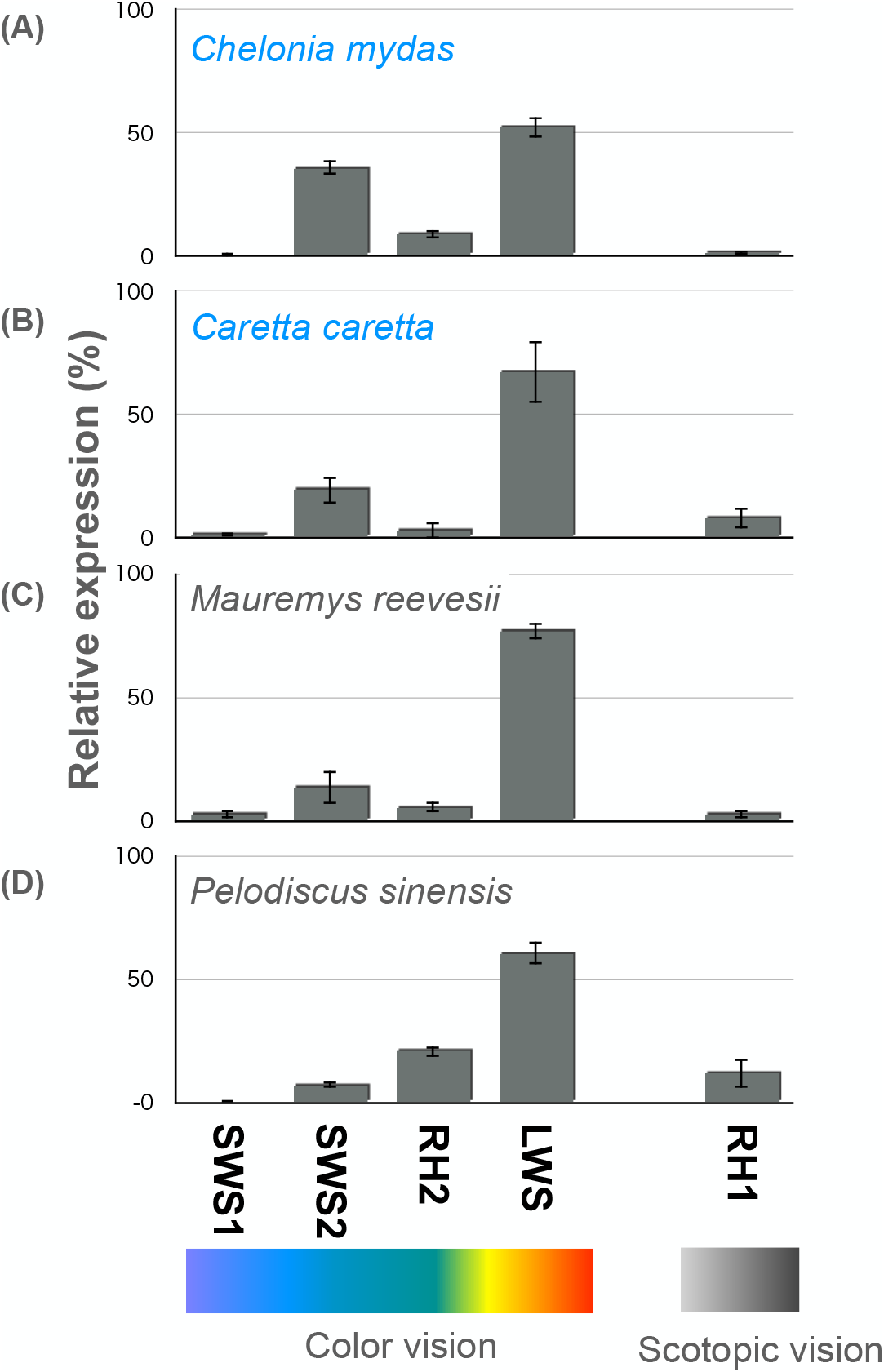
Relative expression of opsin genes in (A) green, (B) loggerhead, (C) three-keeled pond, and (D) softshell turtles.

**Figure 3.**
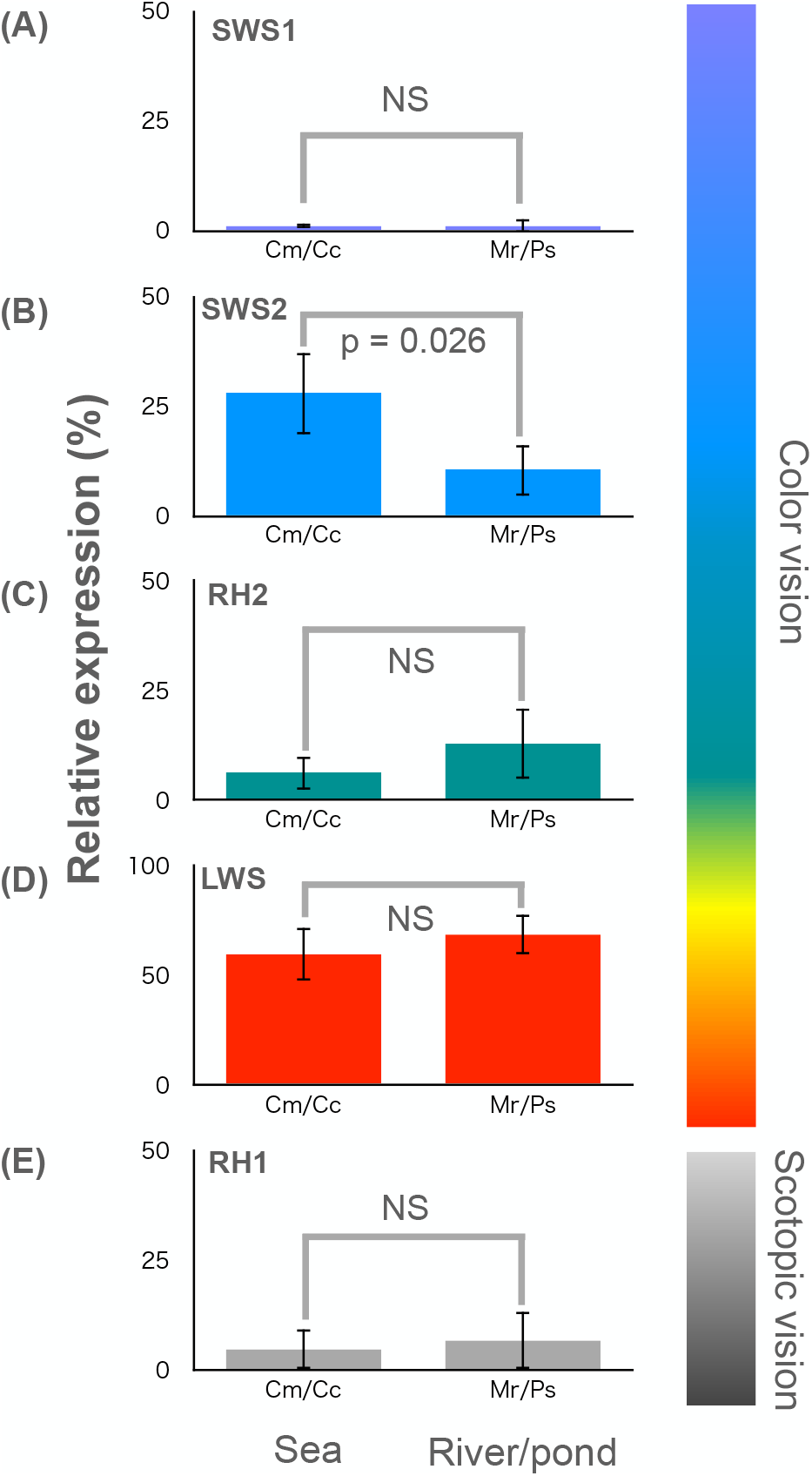
Comparison of the relative expression of (A) SWS1, (B) SWS2, (C) RH2, (D) LWS, and (E) RH1 between sea and freshwater turtles.

## Discussion

In cetaceans and the hydrophiin sea snakes, which are fully adapted to aquatic environments and live their entire lives underwater, the transition from terrestrial to marine environments has affected opsin evolution. Even in pinnipeds and amphibious sea snakes, which are not fully adapted to marine habitats and come ashore to spawn, breed, and rest, the transition to marine environment has significantly affected the sensitivity of opsin pigments. Therefore, we expected the effect of the transition from freshwater to marine environment on opsin evolution in sea turtles.

In previous studies of cetaceans, pinnipeds, and sea snakes, marine adaptation has resulted in shifts in the absorption spectra of opsin pigments and the degeneration or duplication of opsin genes. In sea turtles, however, no gene degeneration, duplication, and amino acid substitutions affecting the sensitivity of opsin pigments have occurred during marine adaptation. Furthermore, we found no amino acid substitution affecting a significant shift in the opsin pigments sensitivity of all turtles analyzed in this study. Most turtles forage in the water and come ashore to bask in the sun. Therefore, they use vision in both terrestrial and underwater light environments. Therefore, opsins are likely conserved due to the functional constraints for using vision in two light environments.

The evolution of opsin expression patterns during marine adaptation has previously been studied in sea snakes, where opsin expression levels tended to be the same in sea snakes and ancestral land snakes (Seiko, et al. 2020). In this study, the expression of *SWS2* was significantly higher in sea turtles than in freshwater ones. The ocean, especially the open ocean, is dominated by blue light (Kirk 1994). Therefore, sea turtles may have adapted their vision to the blue light-rich marine environment by increasing *SWS2* expression.

Previous studies have reported shifts in the absorption of opsin pigments, degeneration, and duplication of opsin genes during marine adaptation. This study shows that the evolution of opsin gene expression may also be part of marine adaptation. Future comparisons of opsin gene expression levels in marine vertebrates and their ancestral species will reveal the evolutionary role of gene expression in marine adaptation.

## Methods

### Samples

Fertilized eggs of the green sea turtle *Chelonia mydas* were collected on Chichijima Island with the permission of the Tokyo Metropolitan Government during the laying periods of 2020. Fertilized eggs of three-keeled pond turtle *Mauremys reevesii* and Chinese softshell turtle *Pelodiscus sinensis* were purchased from turtle farms in Japan, in 2020. Eyeballs were excised from fresh late stage embryos of the three turtle species and used for RNA extraction. Eyeballs were excised from dying juveniles of the loggerhead turtle *Caretta caretta* hatched from the eggs laid by captive females at Kushimoto Marine Park, Japan, in 2020 and used for RNA extraction. All animal experiments were approved by the Committee on the Ethics of Animal Experiments of the Faculty of Science, Toho University (permits 20–51–449). Three individuals were analyzed for each turtle species.

The guidelines for experimental animal management of SOKENDAI were followed throughout the study. The Institutional Animal Care and Use Committee of SOKENDAI approved the animal protocols and procedures.

### RNA extraction, library preparation, and sequencing

Total RNAs were extracted from eyes of green, loggerhead, three-keeled pond, and Chinese softshell turtles using TRIzol RNA Isolation Reagent (Thermo Fisher Scientific, Waltham, MA, USA) according to the manufacturer’s instructions with purification using RNeasy mini kit (Qiagen, Hilden, Germany) following the manufacturer’s protocol. RNA libraries were constructed using the NEBNext Poly(A) mRNA Magnetic Isolation Module and the NEBNext Ultra RNA Library Prep Kit for Illumina (New England Bio Labs, Ipswich, MA) following the manufacturer’s instructions. Short cDNA sequences (paired-end 150 bp) were determined from the libraries using the Illumina HiseqX platform (RNA-seq).

### Phylogenetic analysis

Sequences of the *SWS1, SWS2, RH2, LWS*, and *RH*1 gene of turtles and outgroup species were collected by blastn (Camacho, et al. 2009) from the NCBI nucleotide database using the loggerhead turtle *SWS1, SWS2, RH2, LWS*, and *RH*1 sequences as queries (Table S1). *SWS2* and *LWS* sequences of green turtle and *SWS1* and *SWS2* sequences of softshell turtle were not found in the database. Therefore, we assembled RNA-seq reads and isolated these opsin sequences from each assembled sequence by blastn using the loggerhead turtle opsin sequence as queries. All opsin sequences obtained were aligned with GENETYX ver. 20.0 software (Genetyx Corporation, Tokyo, Japan). The opsin gene trees were constructed by the ML method using the MEGA X software (Kumar, et al. 2018). The model was selected as the best fit model for the ML tree construction using the MEGA X software (Kumar, et al. 2018). The statistical reliability of the tree branches was evaluated using 1,000 bootstrap replicates.

### Expression analysis

RNA-seq reads from each of three individuals of green, loggerhead, three-keeled pond, and softshell turtles were mapped to the sequences of *SWS1, SWS2, RH2, LWS*, and *RH1* from each species. Reads showing similarity (90%) with 90% read lengths were mapped to opsin sequences and expression levels were calculated using CLC Genomics Workbench ver. 11 (https://www.qiagenbioinformatics.com/). TPM (Transcript Per Million) was used for normalized expression values.

