## Supplementary material for "Visual adaptation of opsin gene expression to the aquatic environment in sea turtles": Table S1

Table S1 Sequences used in this study

| | Species | Accession number |
|---|---|---|
| SWS1 | Caretta caretta | XM_048836794.1 |
| | Chelonia mydas | XM_037889438.1 |
| | Dermochelys coriacea | XM_038397580.1 |
| | Trachemys scripta elegans | XM_034763752.1 |
| | Terrapene carolina triunguis | XM_024194545.1 |
| | Chrysemys picta bellii | XM_005281289.2 |
| | Malaclemys terrapin pileata | XM_054011285.1 |
| | Mauremys reevesii | XM_039519978.1 |
| | Mauremys mutica | XM_045030337.1 |
| | Gopherus evgoodei | XM_030582861.1 |
| | Gopherus flavomarginatus | XM_050930114.1 |
| | Chelonoidis abingdonii | XM_032802179.1 |
| outgroup | Varanus komodoensis | XM_044429417.1 |
| SWS2 | Caretta caretta | XM_048833219.1 |
| | Dermochelys coriacea | XM_038381447.2 |
| | Chrysemys picta bellii | XM_005281281.3 |
| | Malaclemys terrapin pileata | XM_054014945.1 |
| | Mauremys mutica | XM_045000692.1 |
| | Mauremys mutica | XM_045000583.1 |
| | Mauremys reevesii | XM_039510324.1 |
| | Gopherus flavomarginatus | XM_050928714.1 |
| | Gopherus flavomarginatus | XM_050927344.1 |
| outgroup | Dromaius novaehollandiae | XM_026094007.1 |
| RH2 | Caretta caretta | XM_048826696.1 |
| | Chelonia mydas | XM_007063507.2 |
| | Dermochelys coriacea | XM_038379721.1 |
| | Trachemys scripta elegans | XM_034767392.1 |
| | Terrapene carolina triunguis | XM_026655132.1 |
| | Terrapene carolina triunguis | XM_024199187.2 |
| | Malaclemys terrapin pileata | XM_054028257.1 |
| | Chrysemys picta bellii | XM_005309675.2 |
| | Mauremys mutica | XM_045013912.1 |
| | Mauremys reevesii | XM_039538204.1 |
| | Chelonoidis abingdonii | XM_032768690.1 |
| | Gopherus evgoodei | XM_030561359.1 |
| | Gopherus flavomarginatus | XM_050953088.1 |
| | Pelodiscus sinensis | XM_006119345.3 |
| outgroup | Falco rusticolus | XM_037410806.1 |
| LWS | Caretta caretta | XM_048832768.1 |
| | Mauremys mutica | XM_045000585.1 |
| | Mauremys reevesii | XM_039511073.1 |
| | Chrysemys picta bellii | XM_005281282.3 |
| | Malaclemys terrapin pileata | XM_054014960.1 |
| | Gopherus flavomarginatus | XM_050928050.1 |
| | Gopherus flavomarginatus | XM_050927343.1 |
| | Dermochelys coriacea | XM_043501319.1 |
| | Pelodiscus sinensis | XM_014581850.2 |
| outgroup | Harpia harpyja | XM_052779277.1 |
| RH1 | Caretta caretta | XM_048858495.1 |
| | Chelonia mydas | XM_007059947.3 |

| | Species | Accession number |
|---|---|---|
| | Dermochelys coriacea | XM_038408468.2 |
| | Trachemys scripta elegans | XM_034777143.1 |
| | Chrysemys picta bellii | XM_008168043.2 |
| | Mauremys reevesii | XM_039547817.1 |
| | Terrapene carolina triunguis | XM_024207554.1 |
| | Mauremys mutica | XM_045024283.1 |
| | Malaclemys terrapin pileata | XM_054036134.1 |
| | Chelonoidis abingdonii | XM_032784259.1 |
| | Gopherus evgoodei | XM_030570330.1 |
| | Gopherus flavomarginatus | XM_050958404.1 |
| | Pelodiscus sinensis | XM_006132837.3 |
| outgroup | Crocodylus porosus | XM_019536553.1 |
